# Mismatch novelty exploration training shifts VPAC_1_ receptor mediated modulation of hippocampal synaptic plasticity by endogenous VIP in male rats

**DOI:** 10.1101/2022.12.21.521348

**Authors:** F Aidil-Carvalho, A Caulino-Rocha, JA Ribeiro, D Cunha-Reis

## Abstract

Novelty influences hippocampal-dependent memory through metaplasticity. Mismatch novelty detection activates the human hippocampal CA1 area and enhances rat hippocampal-dependent learning and exploration. Remarkably, mismatch novelty training (NT) also enhances rodent hippocampal synaptic plasticity. Inhibition of VIP interneurons by prefrontal cortex GABAergic projections promotes rodent exploration. Since VIP, acting on VPAC_1_ receptors (Rs), restrains hippocampal LTP and depotentiation by modulating disinhibition we now investigated the impact of NT on VPAC_1_ modulation of hippocampal synaptic plasticity in male Wistar rats. NT enhanced both CA1 hippocampal LTP and depotentiation unlike exploring an empty holeboard (HT) or a fixed configuration of objects (FT). Blocking VIP VPAC_1_Rs with PG 97-269 (100nM) enhanced both LTP and depotentiation in naïve animals but this effect was less effective in NT rats. Altered endogenous VIP modulation of LTP was absent in animals exposed to the empty environment (HT). HT and FT animals showed mildly enhanced synaptic VPAC_1_R levels but neither VIP nor VPAC_1_R levels were altered in NT animals. Conversely, NT enhanced the GluA1/GluA2 AMPAR ratio and gephyrin synaptic content but not PSD-95 excitatory synaptic marker. In conclusion, NT influences hippocampal synaptic plasticity by reshaping brain circuits modulating disinhibition and its control by VIP-expressing hippocampal interneurons while upregulation of VIP VPAC_1_Rs is associated to maintenance of VIP control of LTP in FT and HT animals. This suggests VIP receptor ligands may be relevant to co-adjuvate cognitive recovery therapies in aging or epilepsy, where LTP/LTD imbalance occurs.

**Significance statement:** This work sheds light into the contribution of hippocampal neuropeptide VIP and VIP containing GABAergic neurons on the reformulation of hippocampal circuit communication leading to altered synaptic plasticity responses upon training in repeated exposure to object mismatch novelty as compared with routine training paradigms with and without objects. Although still far from the clinic, the results suggest that VIP receptor ligands may be useful to co-adjuvate cognitive stimuli therapies based on these aspects of novelty, aiming at the cognitive rescue of aged individuals or epilepsy patients, that show impaired capacity in novelty induced adaptations of cognitive function.

**Graphical abstract:** 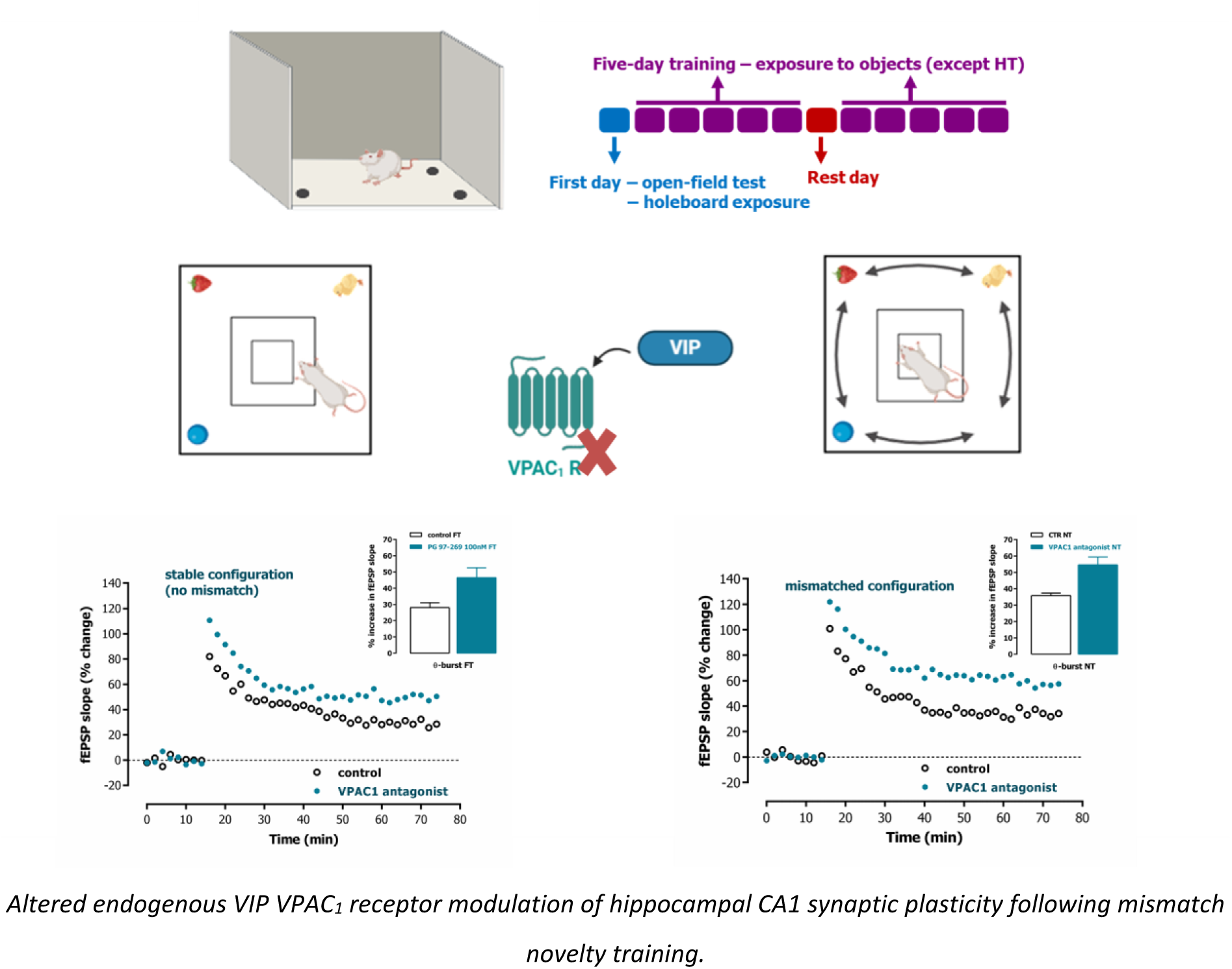

## 1. Introduction

Adaptations in synaptic plasticity events like long-term potentiation (LTP), long-term depression (LTD) or depotentiation of synaptic communication are triggered by recent behavioural experience and contribute to learning and memory and memory stability through metaplasticity (Abraham and Bear, 1996; Izquierdo et al., 2001). While, LTP underlies hippocampal dependent memory acquisition, LTD complements LTP, inducing memory reformulation and reconsolidation, required for lasting memory storage during spatial memory formation (Manahan-Vaughan and Braunewell, 1999; Ge et al., 2010; Alberini and Ledoux, 2013; Dong et al., 2013). In fact, LTD and LTP encode distinct aspects of novelty acquisition, LTD being favoured by exposure to novel objects or novel object location, and LTP being facilitated by exposure to new environments, that, in turn occlude LTD (Kemp and Manahan-Vaughan, 2004). Furthermore, depotentiation mediates reversal of LTP preceding memory reformulation/erasure and may undergo metaplastic control by prolonged or repeated novelty exposure (Qi et al., 2013).

Mismatch novelty, defined by a conflict between expected and perceived temporal sequence or spatial location of items or events, activates the hippocampal CA1 area in human studies and may be a relevant stimulus for cognitive stimulation (Thakral et al., 2015; Trempler et al., 2017). Studies in rodents revealed that when a mismatch occurs between observed and expected scenarios one of two processes is triggered, either memory reconsolidation or memory extinction (Pedreira et al., 2004). Furthermore, mismatched location of known objects was found to enhance inhibitory avoidance learning and memory stability, while favouring hippocampal LTD (Dong et al., 2012). However, how recurrent mismatch novelty detection / exposure affects the hippocampal memory systems in the long term has scarcely been investigated. Recently, we developed a cognitive stimulus program based on mismatch novelty exploration and observed that it induced a long-lasting enhancement of both hippocampal LTP and depotentiation (Aidil-Carvalho et al., 2017), while mildly enhancing hippocampal-dependent learning in young rats (Cunha-Reis, 2020). This suggests that human therapies based on such an approach could be useful for human memory rescue in aging and diseases where LTP/LTD is imbalanced, such as temporal lobe epilepsy (TLE) (Aidil-Carvalho et al., 2017; Cunha-Reis et al., 2021). Vasoactive intestinal peptide (VIP), an important neuromodulator in the central and peripheral nervous system, has also important neurotrophic and neuroprotective actions (Ribeiro et al., 2001; Cunha-Reis et al., 2021). In the hippocampus VIP is only present in GABAergic interneurons (Acsády et al., 1996). and controls hippocampal GABA release, GABAergic transmission and pyramidal cell activity (Cunha-Reis et al., 2017; Cunha-Reis and Caulino-Rocha, 2020). Hippocampal actions of VIP are mediated by activation of two selective receptors, VPAC_1_ and VPAC_2_, but only VPAC_1_ receptors are involved in the inhibition of hippocampal LTP, LTD and depotentiation by endogenous VIP (Cunha-Reis et al., 2014; Caulino-Rocha et al., 2022) that occurs through mechanisms that are fully dependent on GABAergic transmission. Three distinct subtypes of VIP interneurons are present in the hippocampus, hippocampal basket cells directly targeting pyramidal neurons and two interneuron-selective interneuron populations (Acsády et al., 1996), that when activated, may promote hippocampal disinhibition, i.e., inhibition of inhibitory interneurons targeting pyramidal cells. This suggests that different behavioural states may lead to the recruitment of distinct VIP-expressing interneurons with relevant influence on hippocampal synaptic plasticity. In fact, VIP-mediated hippocampal disinhibition was shown to be crucial for goal-directed spatial learning tasks (Turi et al., 2019), and a subpopulation of VIP expressing interneurons recruited during theta oscillations (Luo et al., 2020) may have a role in information gating during spatial navigation memory encoding, while inhibition of VIP interneurons may enhance feedforward inhibition to facilitate hippocampal object representation during spatial exploration (Malik et al., 2022). This is corroborated by the fact that VIP expressing interneurons are targeted by *medium raphe* serotonergic projections as well as septal cholinergic and GABAergic fibres, controlling pacing, engagement, and suppression of hippocampal theta rhythm (Borhegyi et al., 2004; Vandecasteele et al., 2014; Vinogradova et al., 1999). VIP interneurons are also targeted by prefrontal cortex long-range GABAergic projections (Malik et al., 2022), involved in the top-down control of hippocampal function. Their inhibition by these was shown to promote rodent exploration.

In this work, we investigated if the effects of mismatch novelty training on LTP and depotentiation involved an alteration in endogenous VIP VPAC_1_ receptor-mediated modulation of hippocampal synaptic plasticity. We found that while both LTP and depotentiation were enhanced by mismatch novelty training, modulation of LTP and depotentiation by endogenous VPAC_1_ receptor activation was reduced, suggesting that endogenous VIP and mismatch novelty training may share common pathways to enhance synaptic plasticity mechanisms.

## 2. Materials and Methods

Juvenile male Wistar rats (3-4 weeks-old, 120-150g at the beginning of training) were kept at the local Animal House (22 °C), Faculty of Medicine, University of Lisbon, under a 12:12-h light/dark cycle with food and water *ad libitum*. Female rats were not used given that hormonal influences on LTP during the oestrus cycle (Warren et al., 1995; Good et al., 1999) could substantially increase the variability of the training outcomes. Prior to the training procedure, rats were handled for three days. All animal training/testing was performed between 10.a.m. and 3 p.m in a sound attenuated room. Rats were introduced in the behaviour room 30m before the training started. Possible deficits in their motor capacity or altered anxiety levels were evaluated using the open-field (OF) and elevated plus maze (EPM) tests, respectively, to detect deviations that could potentially bias our study. None of the tested animals was excluded from the study since no statistical outliers in the behavioural parameters probed were detected. Animals then underwent a two-week training program in mismatch novelty exploration and its impact on hippocampal synaptic plasticity and its modulation by VPAC_1_ receptors was tested *in vitro* using electrophysiological recordings as previously described (Aidil-Carvalho et al., 2017). VIP and VPAC_1_ receptor levels were evaluated by western blot in hippocampal synaptosomes obtained from the contralateral hippocampus. All practices were in accordance with the Guide for Care and Use of Laboratory Animals, the Portuguese and European law on animal welfare (EU Directive 2010/63/EU for animal experiments) and were approved by the Ethical Committee of the Faculty of Medicine. A preliminary account of some of the results was published as an abstract (Cunha-Reis, 2020).

### 2.1 Open-field (OF) and Elevated Plus Maze (EPM) pre-evaluation tests

The OF test consisted of the exploration of a large square chamber (66cmx66cm wide, 60cm-high wall, Supp. Fig 1.A), for 5 minutes (Crusio et al., 1989). Three virtual zones (central square 20x20cm, intermediate zone and peripheral 15cm-strip adjacent to the walls) were considered for behaviour analysis. Animals were introduced directly into the centre of the apparatus and locomotion in the arena was recorded. Animal performance was evaluated by quantifying the escape latency (s), total distance travelled (cm), the number of rearings (rat standing on its hind legs, playing or not with his front paws on the wall) and the number of entries and the time spent in each virtual zone.

The EPM (Supp. Fig. 1.B) was comprised by two open arms (50cmx10cm) and two enclosed arms (50cmx10cmx40cm), extending from a central platform (10cmx10cm) and raised 50cm above floor level. The animal was positioned on the central platform, facing the open arm, and allowed to explore the maze for 5 minutes. The maze was cleaned with a 70% ethanol solution between trials. Each test was video recorded and later analysed with video-tracking software (Anymaze, Stoelting Europe). The number of entries in open/closed arms, time spent in open/closed arms or the central platform, distance (cm) travelled in the maze, and number of rearings were evaluated (Schneider et al., 2011).

### 2.2 Mismatch novelty training

Animals were randomly assigned to one of three experimental groups. Animals undergoing mismatch novelty training (NT, Fig. 1.A-B) were daily exposed for two weeks to three objects (plastic ball, strawberry-shaped rubber and animal-shaped painted wood figure) always presented in a new location (Aidil-Carvalho et al., 2017) of a holeboard (66x66cm, 60cm-high walled arena containing one hole - 5.5cm diameter, 4.5cm deep-at each corner). As control for novelty training, two animals were exposed to either a fixed object’s distribution (FT, Fig. 1.B), or to the holeboard without any object (HT, Fig. 1.B). Naïve animals were left without training. Training was performed in 5-min sessions once a day. Animals explored the empty holeboard on the first day and objects were introduced in three of the four holes for the NT and FT animals on the second day. FT animals found the objects in this same location in the following days, unlike NT animals. Training was performed in two sets of 5-day consecutive training separated by one resting day (Fig. 1.A). Video monitored trials were recorded as a track file using automated video-tracking software (Smart 2.5, PanLab, Barcelona) for later analysis. For each trial, travelled distance, number of entries and time spent in each virtual zone of the apparatus (central, intermediate, and peripheral zones; Fig. 1.B) were evaluated. The number of nose pokes (head dips into the holes) was used to assess object exploration; the number of rearings and distance travelled taken as a measure of general exploratory activity.

**Figure 1.**
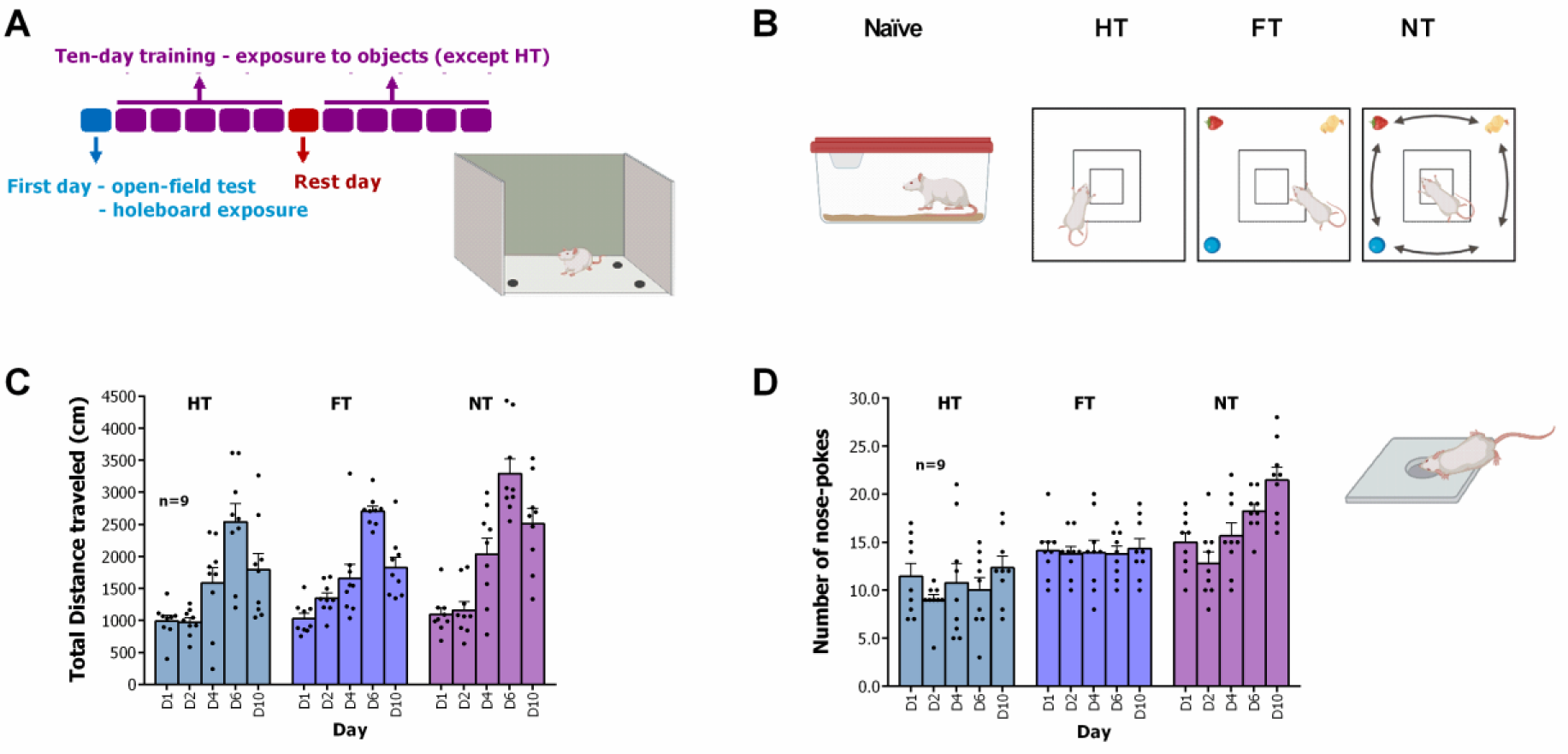
– Mismatch novelty training paradigm and behavioural analysis. **A.** Time-course of the two-week mismatch novelty training and illustration of the holeboard arena used to conduct it. **B.** Mismatch novelty training (NT) paradigm in object exploration in a known environment and respective functional controls: exploration of a fixed configuration of objects (FT) or the empty holeboard (HT, holes not shown). Naïve animals remained every day in their cages. Total distance travelled (**C.**) and number of nose pokes (**D.**) during the training by animals undergoing different training modalities (HT, FT and NT). Individual observations and mean ± S.E.M values are depicted. Differences in the number of nose pokes were significant for FT and NT vs. HT (P<0.01, F_(2,24)_=12.90, Two-way repeated measures ANOVA).

### 2.3 Electrophysiological recordings and synaptic plasticity

Electrophysiological recordings were performed in hippocampal slices of trained rats 3-5 days after training completion as described (Aidil-Carvalho et al., 2017). Hippocampal slices (400μm) cut perpendicularly to the hippocampus long axis were allowed energetic and functional recovery for 1 h at RT in aCSF (mM: NaCl 124, KCl 3, NaH_2_PO_4_ 1.25, NaHCO_3_ 26, MgSO_4_ 1, CaCl_2_ 2, glucose 10) bubbled with 95% O_2_ / 5% CO_2_. Each slice was placed in a recording chamber and superfused (3 ml/min) with gassed aCSF at 30.5°C. Alternate stimulation of two independent sets (S1 and S1) of the Schaffer collateral/commissural fibres was performed every 10 s (rectangular pulses of 0.1 ms) using two bipolar concentric wire electrodes. Field excitatory post-synaptic potentials (fEPSPs) were recorded extracellularly from CA1 *stratum radiatum* using micropipettes (4M NaCl, 2–4 MΩ). Stimulation elicited a fEPSP of 500–850 µV amplitude (about 50% of maximum, similar magnitude for both pathways). The average of six consecutive responses was measured, graphically plotted and recorded for further analysis using the LTP software (Anderson and Collingridge, 2001). fEPSP magnitude was quantified as the slope of the initial phase of the potential.

Independence of the two pathways was confirmed at the end of the experiments by investigating paired-pulse facilitation (PPF) across both pathways. PPF was elicited by two stimuli, with 50 ms pulse interval, delivered sequentially to the two Schaffer collateral pathways (S1 and S2). The P2/P1 ratio between the fEPSP slopes elicited by the second P2 and the first P1 stimuli was computed and compared with the P2/P1 ratio under normal stimulation conditions (10s pulse interval between S1 and S2). Pathway independence was ruled out when PPF of the P2/P1 ratio was larger than 5%.

To elicit LTP, mild theta-burst stimulation (TBS5x4, five trains of 100 Hz, 4 stimuli, separated by 200 ms) was applied. This allowed us to work very far from LTP saturation, that could limit LTP enhancement with concomitant training and addition of the VPAC_1_ receptor antagonist. To obtain depotentiation, LTP was first induced by a slightly more robust theta-burst stimulation (ten trains of 100 Hz, 4 stimuli, separated by 200 ms) and depotentiation was induced 1 h later with low-frequency stimulation (LFS, 1 Hz, 16 min). Stimulation protocols used to induce synaptic plasticity were applied after having a stable baseline for at least 20 min. Stimulus intensity was not changed therein. LTP or depotentiation were quantified as the % change in fEPSP slope 50 to 60 min after the induction protocol, in relation the fEPSP slope measured during the 12 min that preceded it. Changes in the induction phase of LTP or depotentiation were quantified in the 4 to 6 min following the induction protocol. Control and test conditions were tested in independent pathways in the same slice. S1 denotes the pathway (left or right) to which TBS5x4 (or TBS10x4 followed by LFS) was first applied. The VPAC_1_ receptor antagonist, PG 97-269, was added to the perfusion solution 20 min before TBS5x4 or LFS of the test pathway (S2) and kept until the end of the experiment. Each *n* represents a single LTP (or depotentiation) experiment (S1 *vs* S2 conditions) performed in one slice from an independent animal.

### 2.4 Western-blot analysis of VIP and VPAC1 receptor levels, AMPA receptor subunit ratio and GABAergic and glutamatergic synaptic markers

For studying changes in VPAC_1_ receptor expression that could be relevant for synaptic plasticity, we used hippocampal synaptosomes, thus avoiding probing VPAC_1_ receptors in other cell types, like microglia (Cunha-Reis et al., 2021), that could mask small variations occurring in synaptic contacts. Hippocampal synaptosomes were isolated from the hippocampal tissue of trained animals as previously described (Caulino-Rocha et al., 2022). For western blot studies samples were incubated at 95°C for 5 min with Laemmli buffer (125mM Tris-BASE, 4% SDS, 50% glycerol, 0,02% Bromophenol Blue, 5% 1,4-dithiothreitol), run on standard 10% SDS-PAGE and transferred to PVDF membranes (0.45 μm pore, Immobilon) (Rodrigues et al., 2021). These were blocked with either 3% BSA or 5% milk and incubated overnight at 4°C with rabbit polyclonal anti VPAC_1_ receptor (1:600, Alomone Labs Cat# AVR-001; RRID: AB_2341081), rabbit polyclonal anti VIP (1:300, Proteintech, Cat# 16233-1-AP, RRID: AB_2878233), mouse monoclonal anti-gephyrin (1:3000, Synaptic Systems Cat# 147011, RRID:AB_2810215), rabbit polyclonal anti-PSD-95 (1:750, Cell Signalling Technology Cat# 2507, RRID:AB_561221), rabbit polyclonal anti-synaptophysin (1:7500, Synaptic Systems Cat# 101002, RRID:AB_887905), rabbit polyclonal anti-GluA1 (1:4000, Millipore Cat# AB1504; RRID:AB_2113602), rabbit polyclonal anti-GluA2 (1:1000, Proteintech Cat# 11994-1-AP; RRID: AB_2113725), and either mouse monoclonal anti-β-actin (1:5000, Proteintech Cat# 60008-1-Ig, RRID: AB_2289225) or rabbit polyclonal anti-α-tubulin (1:4000, Proteintech Cat# 11224-1-AP, RRID:AB_2210206) primary antibodies. After washing, membranes were incubated for 1h with anti-rabbit or anti-mouse IgG secondary antibody conjugated with horseradish peroxidase (HRP) (Proteintech) at RT. HRP activity was detected by enhanced chemiluminescence using the ImageQuant™ LAS 500 detector (GE Healthcare). Band intensity was evaluated with the Image J software using β-actin or α--tubulin band density as loading control. Band intensities of targeted proteins were normalized to the β-actin or α--tubulin loading controls, respectively.

### 2.5 Materials

PG 97-269, (Phoenix peptides, Europe) was made up in 0.1mM stock solution in CH_3_COOH 1% (v/v). The maximal concentration of CH_3_COOH delivered to the slices, 0.001% (v/v), induced no change on fEPSP slope (n=26). Stock solutions were kept frozen at -20°C in aliquots until use. Aliquots were thawed and diluted in aCSF for use in each experiment.

### 2.6 Statistics

Values are presented as the mean ± S.E.M and n denotes the number of animals in all Figures. We used one slice per animal and condition in all electrophysiological experiments. A total of 35 animals were used in this study, 8 control, and 9 for each trained group. Number of animals was the necessary to obtain statistical significance in synaptic plasticity experiments (from 5-7, as inferred from previous electrophysiology experience) and to obtain required tissue for WB experiments. Differences in behavioural parameters for the EPM and OF tests were analysed by One-way ANOVA (using Sidaks’s correction for multiple comparisons). Throughout training significance of the differences in behavioural parameters between the groups (Fig. 1.D-E) was calculated by Two-way repeated measures ANOVA (using Tukey’s correction for multiple comparisons). Significance of potentiation or depotentiation against the null hypothesis was evaluated using Student’s t-test. Significance of differences in LTP between control and PG 97-269 obtained in the same slices were calculated using paired Student’s t-test. Significance of differences caused by different modes of training on LTP, depotentiation and protein levels as determined by western blot were calculated by One-way ANOVA. No outliers were identified in our data (ROUT method). All statistical analysis was performed using GraphPad Prism 6.01. P<0.05 or less denotes significant differences.

## 3. Results

### 3.1 Open-field (OF) and Elevated Plus Maze (EPM) tests

Analysis of behaviour in the OF, relied on the definition of three virtual zones (peripheral, intermediate, and central zones). Animals spent much less time in the centre (3.7±0.9s, n=26) and intermediate (6.0±1.2s, n=26) zones of the apparatus when compared to the periphery zone (286.7±2.0s, n=26), a behaviour known as thigmotaxis (F _(2,75)_ =12838, P<0.0001, Supp. Fig. 1.A). Accordingly, the distance travelled in the centre and intermediate zones (72.6±13.2cm and 123.1±23.5cm, n=26, respectively) of the apparatus was much smaller (F _(2,75)_ =144.9, P<0.0001) when compared to the periphery zone (3174.0±254.2cm, n=26). Animals had very few entries (1-4) in any of the different zones. Global locomotion (accessed by total distance travelled; 3460.2±219.0cm, n=26) was high as usually observed in our lab for juvenile rats. The total number of rearings (9.2±1.6, n=26) reflects mostly rearings performed in the periphery zone.

In the EPM (Supp. Fig. 1.B) the %time animals spent in the open arms (19.2±2.3% of a 5-min trial, n=26) was much lower (F _(2,75)_ =124.4, P<0.0001) than the %time spent in the closed arms (63.3±3.0%, n=26). In the remaining time (17.6±1.3%, n=26) the animals remained in the centre of apparatus (the crossing of closed and open arms). Accordingly, all animals entered the open arms less (5.3±0.5, n=26) than the closed arms (8.6±0.4, n=26) from this central position. Rearings (7.3±0.8, n=26) were almost entirely performed within the closed arms.

### 3.2 Animal training in novel location of known objects

The two-week NT program (Aidil-Carvalho et al., 2017) consisted 5-min per day training in the novel location of known objects delivered in two series of 5-day training. As a control, animals were exposed either to the holeboard in the absence of objects (HT) or to the known objects every day in a fixed location (FT) of the holeboard. Behaviour was observed throughout the training sessions.

Upon initial exposure to the holeboard (no objects for all animal groups), the general exploratory behaviour was reduced when compared to the one observed in the OF maze. The number of rearings in holeboard exploration (5.3±0.3, n=27), was less than the one observed in the open-field test (9.2±1.6, n=26) and the total distance travelled on the holeboard (2273.8±201.6cm, n=27) was slightly higher than the total distance travelled in the open-field (2252.9±96.0cm, n=26). The animals spent more time in the periphery zone of the holeboard (276.9±5.2s, n=27), where the holes were located, than in the periphery of the OF (233.1±8.0s, n=26). Thus, animals although likely more familiar with the arena, did not spend more time in the intermediate or centre zones, remaining in the periphery exploring the holes, as evidenced by the number of nose-pokes (12.6±0.8, n=27). The next day, when objects were placed in the holes for NT and FT animals, the number of nose-pokes increased to17.0±1.0 (n=9) and 16.3±1.2 (n=9), respectively (F _(2,24)_ = 6.91; P<0.01). This was not observed for the HT group (11.3±1.3, n=9). No significant differences were encountered in the distance travelled in the holeboard periphery zone as compared with the first exposure to the empty holeboard between for any of the trained groups. The number of rearings decreased mildly for all animal groups, likely due to a diversion of their focus to exploration of the holes instead of the whole holeboard.

A marked enhancement in the total distance travelled was observed with training day for all animals (Figure 1.C, F_(4,96)_ = 59.0, P<0.0001, two-way repeated measures ANOVA). Total distance travelled significantly higher for NT vs. HT animals (P<0.05, F _(2,24)_ = 4.20, two-way repeated measures ANOVA). No interaction between the two variables (training mode and training day) was detected (P>0.05, F _(8,72)_ = 0.779, two-way repeated measures ANOVA). The time spent in the periphery of the holeboard varied mildly with training day (F_(4,96)_ = 2.50, P<0.05, two-way repeated measures ANOVA) but not with animal experimental groups (P>0.05, F _(2,24)_ = 0.196, two-way repeated measures ANOVA). No interaction between the two variables (training mode and training day) was detected (P>0.05, F _(8,96)_ = 0.412, two-way repeated measures ANOVA).

The number of nose-pokes varied significantly with daily training (Figure 1.D, F _(4,96)_ = 7.80, two-way repeated measures ANOVA, P<0.01) and with animal training configuration (P<0.01, F _(2,24)_ = 12.90, two-way repeated measures ANOVA). Nose-pokes increased from 15.0±1.0 (n=9) on the first day to 21.4±1.3 (n=9) on the tenth day of training for the NT group. Variation in the number of nose pokes for the FT group (14.1±0.9 on day 1 vs. 14.3±1.0 on day 10, n=9) and for HT 3group (11.4±1.3 on day 1 vs. 12.3±1.2 on day 10, n=9) were not so pronounced. Interaction between the two variables (training mode and training day) was significant (P<0.05, F _(8,96)_ = 7.80, two-way repeated measures ANOVA). The number of rearings increased significantly with daily training in HT but not NT or FT animals (F _(4,96)_= 2.76, two-way repeated measures ANOVA, P>0.05) and increased mildly but significantly for HT vs. FT and (P<0.01, F _(2,24)_ = 8.64, two-way repeated measures ANOVA). No interaction between the two variables (training mode and training day) was detected (P>0.05, F _(8,96)_ = 1.33, two-way repeated measures ANOVA).

### 3.3 Impact of mismatch novelty training on hippocampal synaptic plasticity

Basal stimulation conditions elicited a fEPSP that was 40-60% of the maximal response in each slice and had an average slope of 0.589±0.032mV/ms (n=40). For synaptic plasticity experiments, when inducing LTP, mild theta-burst stimulation (TBS5x4) was applied to one of the pathways (S1 or S2) that was therein considered the control pathway. In Naïve animals an initial post-tetanic potentiation (PTP, 53.5±2.8% increase in fEPSP slope, n=7, Fig. 2.A) was observed that decayed over time. An LTP was still observed 50-60min after stimulation, as a 24.6±2.3% increase in fEPSP slope (n=7, P<0.05, t-test, Fig. 2.A). This LTP was fully dependent on NMDA receptor activity, as previously described by others, since it was abolished (P>0.05, t-test, n=4) when the NMDA receptor antagonist AP-5 (100μM) was present in the medium from 20 min before TBS5x4 stimulation. As previously described, when the VPAC_1_ receptor antagonist was added to the slices 20 min before TBS5x4 delivery to the test pathway, PTP was higher than in the control pathway (81.5±9.5% increase in fEPSP slope) and the observed LTP 50-60 min post stimulation was also enhanced (P<0.05, paired t-test) as it induced a 45.5±6.5% increase (n=7, Fig. 2.A) in fEPSP slope, as previously described (Caulino-Rocha et al., 2022), nearly doubling the potentiation obtained. PG 97-269 (100nM), when added to the slices, did not change the fEPSP slope as estimated in the control pathway (not stimulated with TBS).

**Figure 2.**
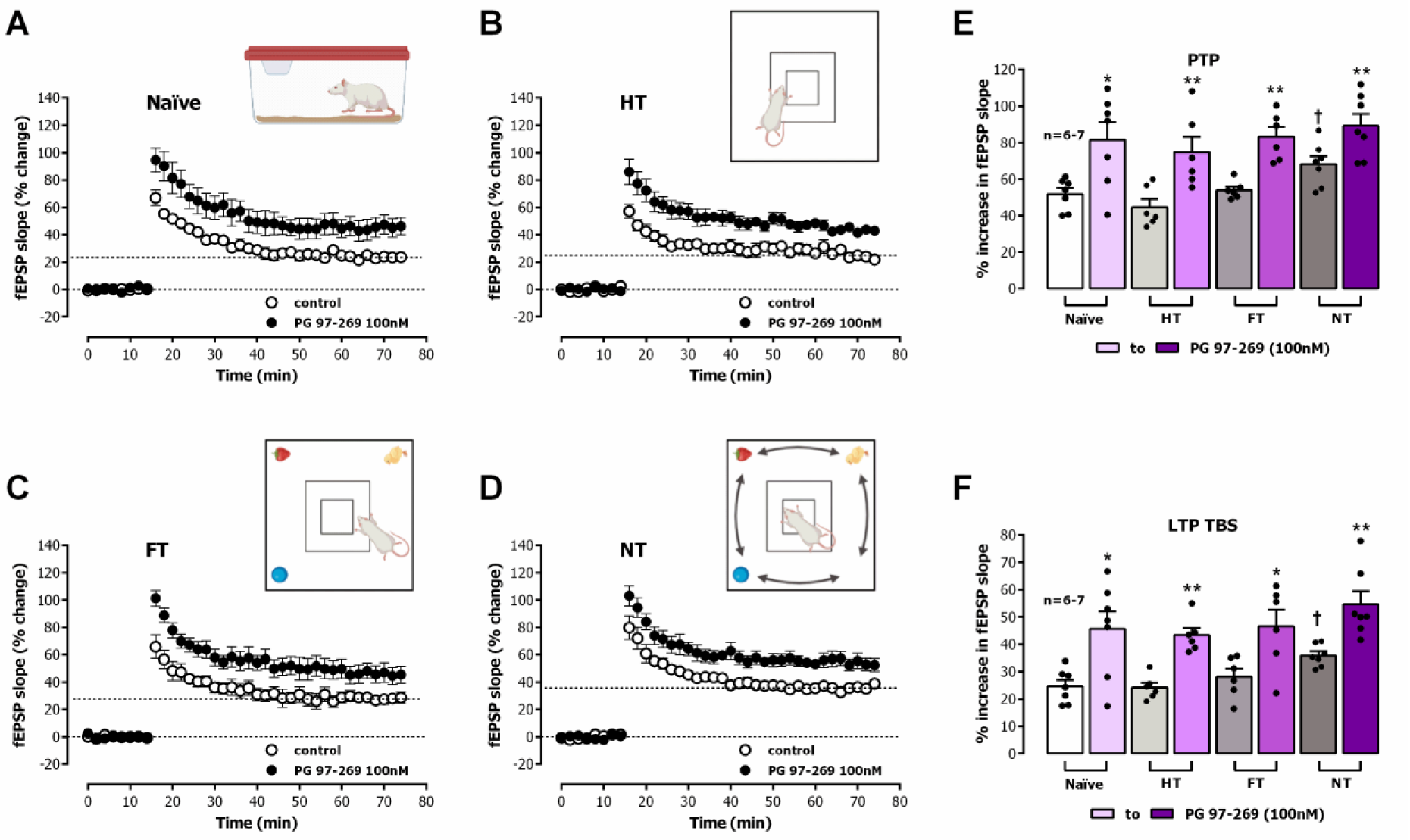
– Impact of mismatch novelty training on VPAC_1_ modulation of TBS-induced CA1 LTP by endogenous VIP. Averaged time-course of changes in fEPSP slope caused by theta-burst stimulation (5x 100 Hz bursts, 200ms interval, ***TBS***, 5x4) in hippocampal slices from Naïve (**A.**), holeboard trained (HT, **B.**), fixed object trained (FT, **C.**) and mismatch novelty trained (NT, **D.**) animals in the absence (-o-) and in the presence (-•-) of the VPAC_1_ receptor antagonist PG 97-269 (100nM). Values are the mean ± S.E.M of 6-7 experiments performed in different individuals. Initial (post-tetanic) potentiation (PTP, **E.**) and magnitude of the LTP (**F.**) estimated from the averaged increase in fEPSP slope observed 4-6 and 50-60 min after ***TBS*** in control conditions for Naïve animals (open bar) and left to right HT (light grey), FT (median grey) and NT (dark grey) animals, and respective PTP (**E.**) and LTP magnitude (**F.**) obtained in the presence of PG 97-269 (100nM) in Naïve animals (light pink) and, left to right, HT (lavender), FT (lilac) and NT (violet) animals. Individual observations and mean ± S.E.M values are depicted. * p < 0.05, ** p < 0.01 and *** p<0.005 (paired Student’s) as compared to PTP (**E.**) or LTP (**F.**) obtained in the absence of PG 97-269 for each training modality (next column in the left). ⩾ p < 0.01 (one-way ANOVA) as compared to naïve animals in the absence of PG 97-269. ‡ p < 0.01 (one-way ANOVA) as compared to naïve animals in the presence of PG 97-269.

As previously described, when TBS5x4 was applied to hippocampal slices obtained from NT rats, an enhanced LTP (F _(3,22)_ =6.22, P=0.004) was observed 50-60min post stimulation, with TBS5x4 causing a 35.8±1.5% increase in fEPSP slope (n=7, Fig. 2.D) in NT animals. Yet, such an enhancement was absent in slices from HT or FT animals, since we encountered a similar LTP magnitude to the one observed in Naïve rats (24.2±1.8% n=6, Fig. 2.B and of 27.8±2.1% n=6, Fig. 2.C enhancement in fEPSP slope for HT and FT, respectively, P>0.05). This difference in the response to TBS5x4 was evident early after LTP induction since 4-6 min post stimulation with TBS5x4 the PTP in NT animals was significantly higher (68.2±4.5% increase in fEPSP slope) than the one observed in Naïve and HT animals (51.7±3.3% and 44.6±4.5% increase in fEPSP slope, respectively).

When PG 97-269 (100nM) was added to the slices 20 min before stimulation with TBS5x4 in the test pathway, the enhancement induced by PG 97-269 on LTP in HT and FT animals was similar (F _(3,22)_ =0.89, P>0.05) to the one observed in Naïve animals (43.3±2.6% n=6, Fig. 2.B and of 46.5±6.1% n=6, Fig. 2.C, enhancement in fEPSP slope for HT and FT, respectively). As in naïve animals, this LTP nearly doubled the effects obtained in the absence of the VPAC_1_ antagonist (P<0.01 and P<0.05, paired t-test). Conversely, endogenous VIP modulation of hippocampal LTP, although still present, was impaired by mismatch novelty training since LTP elicited by TBS5x4 in the presence of 100nM PG 97-269 (54.6±4.8% increase in fEPSP slope, n=7, Fig.2.D) was only about 50% larger (P<0.01, paired t-test) than the one observed in the absence of the VPAC_1_ antagonist in the same slices in NT animals. This effect was even more pronounced in the early moments post-induction, since no significant effect (P>0.05, paired t-test) was observed in NT animals between the PTP obtained in the control pathway (absence of drugs, % increase in fEPSP slope 68.2±4.5, n=7) and the one obtained in the test pathway (presence of PG 97-269 100nM, % increase in fEPSP slope 89.2±6.5, n=7).

Low-frequency stimulation (LFS) delivered to hippocampal slices of adult rats *in vitro* does not produce an LTD of CA1 synaptic transmission in the absence of added drugs (Wagner and Alger, 1995; Cunha-Reis et al., 2014). As such, to induce long-lasting depression of synaptic transmission LFS was applied 1h after delivering a TBS10x4 stimulus. In Naïve animals, a depotentiation of synaptic transmission was immediately observed after LFS stimulation (% decrease in fEPSP slope 20.4±1.8%), that was partially maintained 50-60min after (12.2±1.4% decrease in fEPSP slope, n=5, P<0.05, Fig. 3.A). As previously described (Cunha-Reis et al., 2014), when LFS was delivered in the same slices in the presence of the VIP VPAC_1_ receptor antagonist PG 97-269 (100nM) the depotentiation was about twice as large (26.2±1.0% decrease in fEPSP slope, n=5, Fig. 3.A, P<0.01, paired t-test).

**Figure 3.**
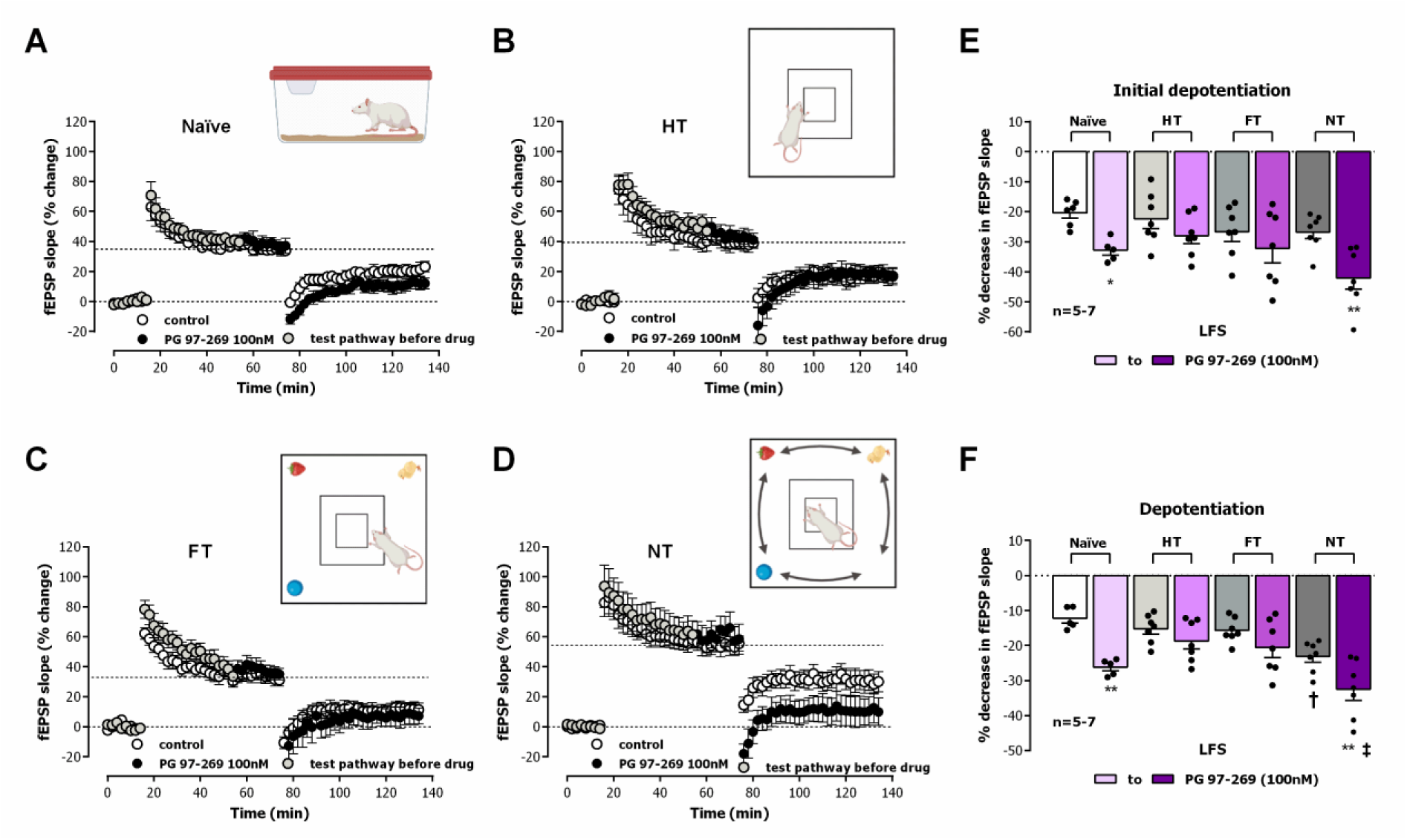
– Impact of mismatch novelty training on VPAC_1_ modulation of CA1 depotentiation by endogenous VIP. Averaged time-course of changes in fEPSP slope caused TBS10x4 followed 1 hour later by LFS (1Hz, 15min) in hippocampal slices from Naïve (**A.**), holeboard trained (HT, **B.**), fixed object trained (FT, **C.**) and novelty trained (NT, **D.**) animals in the absence (-o-) and in the presence (-•-) of the VPAC_1_ receptor antagonist PG 97-269 (100nM) from 20 min before LFS. Dots in the test pathway before addition of PG 97-269 (potentiation curve) are shown in grey. Values are mean ± S.E.M. Initial depression (**E.**) and depotentiation magnitude (**F.**) of the fEPSP slope observed 4-6 and 50-60 min after LFS for Naïve animals (open bar) and left to right HT (light grey), FT (median grey) and NT (dark grey) animals, and respective initial depression (**E.**) and depotentiation magnitude (**F.**) obtained in the presence of PG 97-269 (100nM) in Naïve animals (light pink) and left to right HT (lavender), FT (lilac) and NT (violet) animals. Individual observations and mean ± S.E.M values are depicted. * p < 0.05 and ** p < 0.01 and *** p<0.005 (paired Student’s) as compared to initial depotentiation (**E.**) or depotentiation (**F.**) obtained in the absence of PG 97-269 for each training modality (next column in the left). ⩾ p < 0.01 (one-way ANOVA) as compared to naïve animals in the absence of PG 97-269. ‡ p < 0.01 (one-way ANOVA) as compared to naïve animals in the presence of PG 97-269.

When the same stimulation pattern was delivered to hippocampal slices obtained from NT animals (Fig. 3.D) a larger depotentiation was observed (F _(3;24)_ =8.90; P=0.0003), since LFS delivered 1h after TBS10x4 now caused a decrease of 23.1±1.7% (n=7) in fEPSP slope. This enhancement was not observed for HT and FT animals, since depotentiation magnitude was comparable to the one obtained in Naïve animals 50-60min after LFS (% decrease in fEPSP slope of 15.3±1.5% n=7, Fig. 3.B and of 15.8±1.5% n=7, Fig. 3.C, F _(3;24)_ =8.90; p=0.4248 and 0.3202, respectively). This difference in the response to LFS was not evident early after LTD induction since 5 min post stimulation the depression in fEPSP slope in NT animals was not significantly higher than the one observed in Naïve, HT and FT animals (F _(3,24)_ =1.61, P>0.05, Fig. 3.E).

When PG 97-269 (100nM) was added to the slices 20 min before LFS of the test pathway, the depotentiation in HT and FT was not significantly altered, being only slightly higher than the one observed in the absence of PG 97-269 (18.8±2.2%, n=7, Fig. 3.B and of 20.6±2.9%, n=7, Fig. 3.C, decrease in fEPSP slope for HT and FT, respectively, F _(3,24)_ =1.27, P>0.05). Endogenous VIP modulation of hippocampal CA1 depotentiation was impaired, but still present, in animals undergoing mismatch novelty training (F_(7,44)_ =7.661, P<0.0001) since depotentiation elicited by LFS in the presence of PG 97-269 (100nM) was only about 40% larger (32.6±3.1% increase in fEPSP slope, n=7, Fig.3.D) than the one observed in the absence of the VPAC_1_ antagonist for NT animals. In summary, endogenous VIP modulation of depotentiation was abolished for HT and FT animal training and impaired in NT animals, even though depotentiation itself is enhanced in NT animals.

### 3.4 Impact of mismatch novelty training on hippocampal levels of VIP and synaptic VIP VPAC1 receptors

To determine in differences in VPAC_1_ modulation of hippocampal synaptic plasticity in the hippocampus could be due to training-induced adaptations in VPAC_1_ receptor levels we performed western blot experiments in hippocampal synaptosomes obtained from the hippocampus of NT trained rats and respective naïve and functional training controls (HT and FT animals). VPAC_1_ receptor levels nearly doubled in HT rats since we observed a 72.9±13.6% increase in VPAC_1_ receptor immunoreactivity (n=6, Fig 4, F _(3,20)_ =16.34, P<0.0001, Tukey’s post-hoc test). We also observed a very mild increase (27.9±7.6%, n=6, Fig 4) in VPAC_1_ receptor levels in FT animals but VPAC_1_ receptor levels were not significantly altered in NT rats (n=6, Fig. 4, P>0.05, Tukey’s post-hoc test), thus suggesting that modified control of synaptic plasticity in NT animals is not due to altered VPAC_1_ receptor levels. To determine if altered endogenous VIP levels in hippocampal nerve terminals could account for this effect, we also performed WB experiments to quantify VIP synaptic levels. VIP was mildly enhanced in synaptosomes obtained from HT animals (23.1±6.7%, n=5, Fig 4, F _(3,16)_ =7.45, P<0.01) but not in FT or NT animals, for which a mild but not significant decrease in VIP levels was observed. None of these were significantly different when compared to Naïve animals (n=5, P>0.05, Fig. 4.D), but a significant difference was observed between HT and FT or NT animals (n=5, Fig. 4.D, P<0.01, Tukey’s post-hoc test).

**Figure 4.**
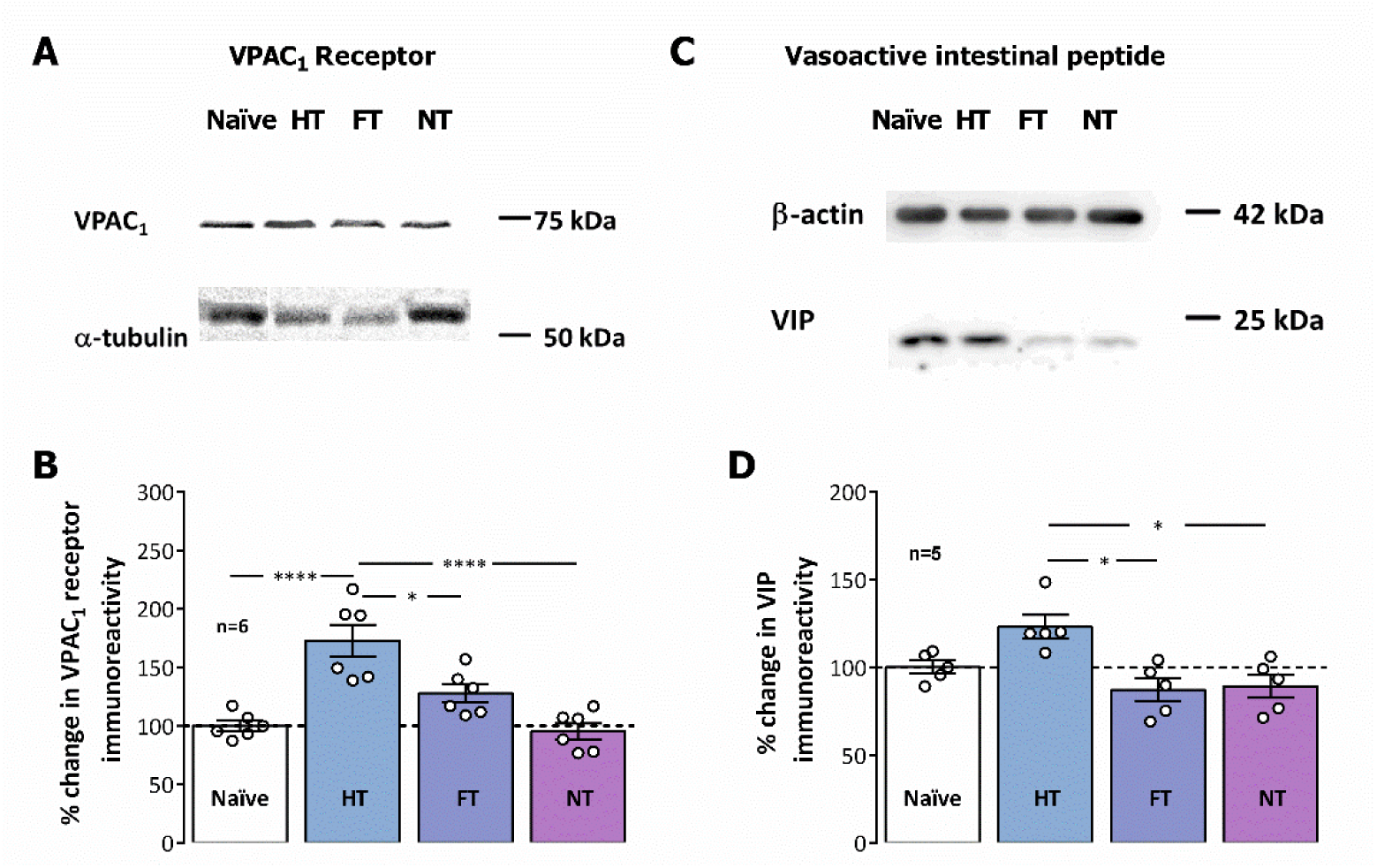
– Impact of mismatch novelty training on VIP and VIP VPAC_1_ receptor levels in hippocampal synaptosomes. Individual experiment showing western-blot immunodetection of VIP VPAC_1_ receptors (**A.**) and respective α-tubulin loading control or VIP and respective β-actin loading control (**C.**). **B.** Increased levels of VPAC_1_ receptors (**B.**) and mildly increased VIP levels (**D.**) observed in HT rats as shown by Western blot immunodetection in hippocampal synaptosomes. Individual observations and mean ± S.E.M values of 5-6 independent experiments performed in duplicate are depicted. 100% - averaged target immunoreactivity obtained for Naïve rats. **\***p < 0.05 and **\*\***p < 0.01 (One-way ANOVA followed by Tukey’s multiple comparison test).

### 3.5 Impact of mismatch novelty training on hippocampal levels of GABAergic and glutamatergic synaptic markers and AMPA receptor subunit ratio

To investigate the impact of NT on hippocampal circuits relevant for modulation of synaptic plasticity we probed the hippocampal levels of synaptic proteins involved in both GABAergic and glutamatergic synaptic transmission, gephyrin and PSD-95, respectively, and a general synaptic marker, synaptophysin. In hippocampal synaptosomes obtained from NT and respective naïve and functional training controls (HT and FT animals), we observed that the synaptic gephyrin levels nearly doubled in HT and FT rats (89.2±14.0% and 84.9±10.2% increase in gephyrin immunoreactivity, n=5, Fig 5.A, F_(3,16)_ =9.820, P<0.005, Tukey’s post-hoc test). This increase in gephyrin levels was tendentially milder (66.6±15.2%, n=5, Fig 5.A, P<0.05, Tukey’s post-hoc test) in NT animals. Altogether, this suggests that modified control of synaptic plasticity in NT animals is influenced by altered GABAergic synaptic function, but it is not clear if this sustains the differences observed in VIP regulation of synaptic plasticity when compared to HT and FT controls. The synaptic levels of PSD-95, a marker of the glutamatergic postsynaptic component, and of synaptophysin, a protein present in synaptic vesicles used general as marker of synapses, were not significantly changed by NT or any of the functional training paradigms used as controls (P>0.05, F _(3,16)_ =2.488 and 1.291, respectively).

**Figure 5.**
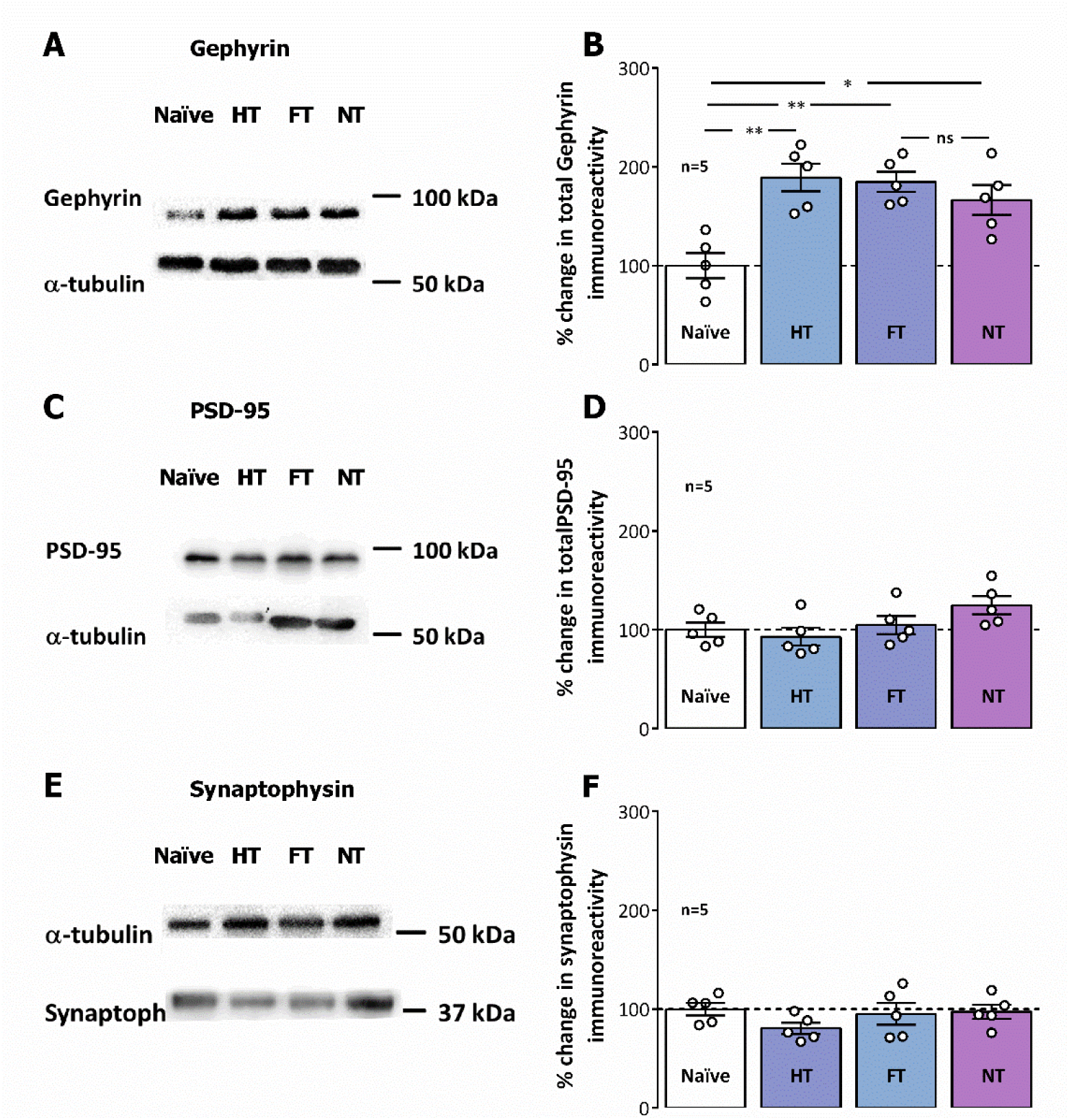
– Impact of mismatch novelty training on GABAergic and glutamatergic markers in hippocampal synaptosomes. Individual experiment showing western-blot immunodetection of gephyrin (**A.**) and respective α-tubulin loading control, PSD-95 (**C.**) and respective α-tubulin loading control, or synaptophysin and respective α-tubulin loading control (**E.**). **B.** Increased levels of gephyrin (**B.**) and undisturbed levels of PSD-95 (**D.**) and synaptophysin (**F.**) observed in HT, FT and NT rats as shown by Western blot immunodetection in hippocampal synaptosomes. Individual observations and mean ± S.E.M values of 5 independent experiments performed in duplicate are depicted. 100% - averaged target immunoreactivity obtained for Naïve rats. **\***p < 0.05 and **\*\***p < 0.01 (One-way ANOVA followed by Tukey’s multiple comparison test).

Given that AMPA receptor subunit composition, a variable arrangement of GluA1-4 subunits, can influence channel function and LTP outcomes (Chater and Goda, 2022) and that AMPA receptors lacking GluA2 are Ca^2+^ permeable, being able to contribute to the postsynaptic Ca^2+^ rise and potentiation levels upon LTP induction, we investigated possible changes in synaptic AMPA receptor subunit composition, specifically synaptic GluA1 and GluA2 levels and the GluA1/GluA2 ratio, that may contribute to altered LTP following NT. In all trained groups (HT, FT and NT animals), the synaptic levels of AMPA GluA1 subunits were mildly enhanced (27-38%, n=5, P<0.05, F _(3,16)_ =3.275, Fig. 6.A). GluA2 subunit levels showed a more prominent enhancement upon HT and FT training (69.3±10.7% and 51.4±12.8%, respectively, n=5, Fig. 6.B, F _(3,16)_ =8.801, P<0.001) yet this enhancement was not present in NT animals (n=5, Fig. 6.B, P>0.05, Tukey’s post-hoc test). As a consequence, the GluA1/GluA2 ratio was mildly higher in NT animals (30.5±7.8%, n=5, P<0.05, F _(3,16)_ =4.850, Fig. 6.C), but not significantly changed (P>0.05, Tukey’s post-hoc test) in HT or FT animals when compared with naïve controls.

**Figure 6.**
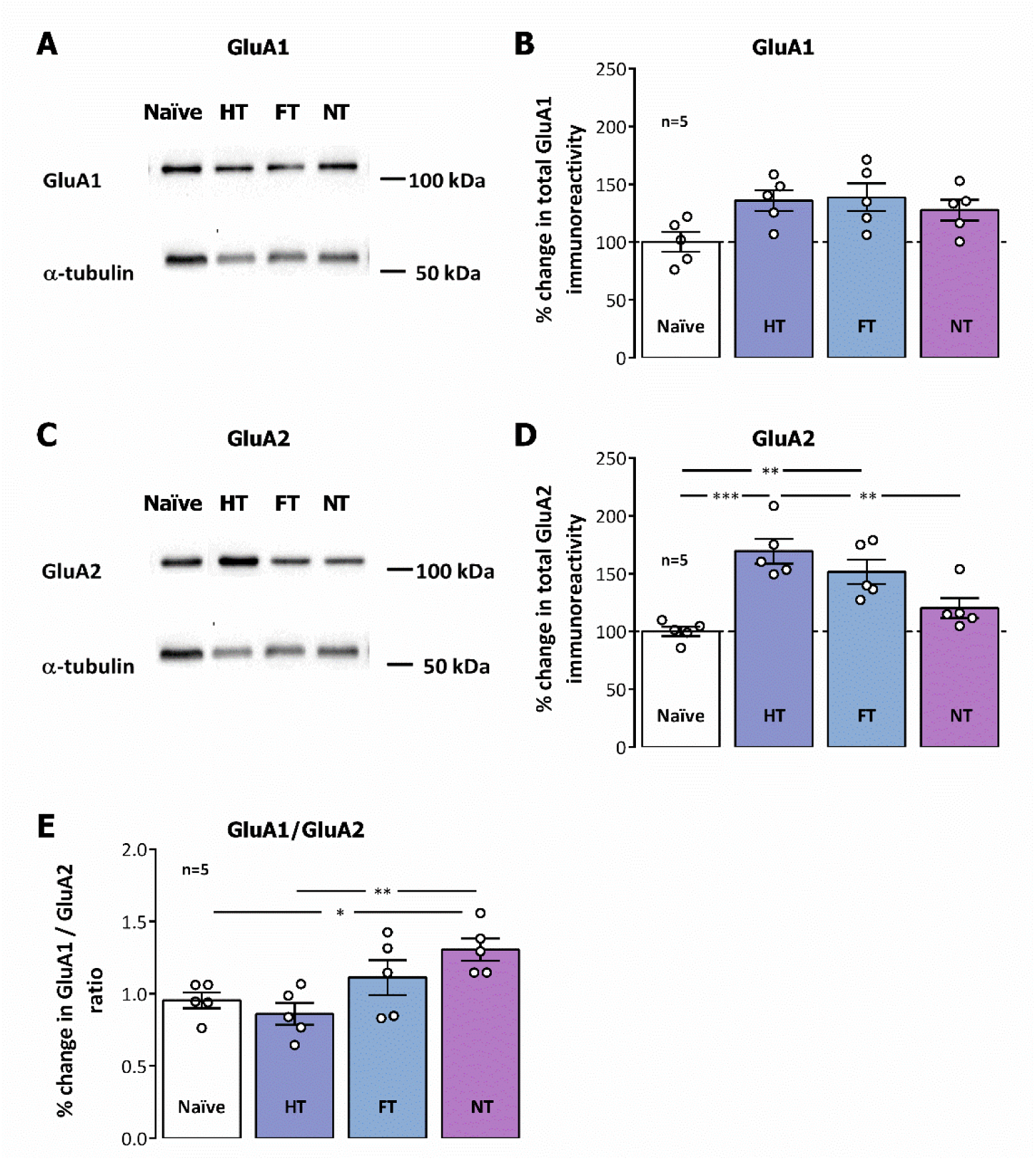
– Impact of mismatch novelty training on AMPA GluA1 and GluA2 levels and GluA1/GluA2 ratio in hippocampal synaptosomes. Individual experiment showing western-blot immunodetection of GluA1 (**A.**), GluA2 (**C.**) and respective α-tubulin loading controls. **B.** Mildly increased levels of GluA1 (**B.**) and substantially increased levels of GluA2 (**D.**) observed in HT, FT and NT rats as shown by Western blot immunodetection in hippocampal synaptosomes. **E.** Increased GluA1/GluA2 ratio observed in NT but not HT and FT animals as compared with naïve controls. Individual observations and mean ± S.E.M values of 5 independent experiments performed in duplicate are depicted. **\***p < 0.05 and **\*\***p < 0.01 (One-way ANOVA followed by Tukey’s multiple comparison test).

## 4. Discussion

The main findings of the present work are that training of young adult Wistar rats in exploration of the novel location of known objects in a holeboard for two weeks induces an impairment in the ability of endogenous VIP to inhibit LTP and depotentiation in hippocampal CA1 area, through activation of VPAC_1_ receptors. This shift does not appear to involve substantially altered synaptic VIP or VPAC_1_ receptor levels in novelty trained rats and is not likely due to a ceiling effect on LTP expression, given the mild TBS train used to induce LTP in this work (Rodrigues et al., 2021). As such, these observations suggest that altered control of hippocampal synaptic plasticity by VIP VPAC_1_ receptors following mismatch novelty training is instead related to alterations in either the targets of VIP immunoreactive interneurons, or global hippocampal network activity. Nevertheless, the HT paradigm used as control caused a mild enhancement of VIP in the hippocampus and a considerable increase in VPAC_1_ receptor levels. Although this could suggest a potential role in enhanced inhibition of synaptic plasticity events in the hippocampus, such an effect was not observed in hippocampal CA1 synaptic plasticity, albeit such altered VPAC_1_ levels may influence synaptic plasticity outcomes in other hippocampal areas not probed in this study. Our findings provide a first insight on a possible mechanism for the selective enhancement of hippocampal synaptic plasticity by mismatch novelty training, the regulation of hippocampal circuits also targeted by interneurons expressing VIP. This could be useful to develop combination therapies (cognitive together with pharmacological) for cognitive decline in aging or epilepsy.

Different aspects of novelty presentation during exploration of the environment have been described to influence cognition in different ways (Izquierdo et al., 2001; Dong et al., 2012; Quent et al., 2021). Mismatch novelty detection is vital for adaptation to novel situations in an otherwise familiar environment or to detect inconsistencies with previously acquired memories. As such, it is not surprising that it primes the brain for cognitive flexibility and learning-associated plasticity (Moncada and Viola, 2007; Park et al., 2021; Quent et al., 2021). This makes it the ideal cognitive training paradigm to elicit changes in synaptic plasticity and cognition, but its underlying mechanisms need to be further elucidated.

In this work, when initiating the training, the rats were first exposed to the empty holeboard. This triggers a behavioural response to the novel environment that was previously shown to reflect a combination of interest in exploration and fear or anxiety (Kanari et al., 2005). We observed that not only general exploratory activity, as assessed by the number of rearings, was decreased when compared to exposure to the OF but also that locomotion in the periphery was slightly enhanced. This likely reflects also an interest in the holes, as was previously evidenced by the increase in the number of nose-pokes (Brown and Nemes, 2008; Aidil-Carvalho et al., 2017). The presence of objects in the holes on the second day of training for NT and FT animals constituted an added novelty and, as expected, induced an enhancement in the number of nose-pokes for both the NT and FT groups but not the HT group, re-exposed to the holeboard, suggesting that the presence of the objects is driving this increase in head-dipping (Aidil-Carvalho et al., 2017). The distance travelled in the periphery (but not the intermediate and central zones) of the holeboard increased along training for NT animals (Fig 1), indicating an enhanced interest in object/hole exploration over holeboard space exploration. Conversely, HT and FT animals demonstrated an increase in general exploration, estimated from the time spent in the intermediate and central zones and number of rearings, confirming that novel object configurations drive enhanced interest in object exploration (Goh and Manahan-Vaughan, 2013b; Aidil-Carvalho et al., 2017).

Mismatch novelty training induced an enhanced LTP response to TBS5x4 stimulation and enhanced depotentiation response to LFS in NT animals but not HT or FT animals as previously shown (Aidil-Carvalho et al., 2017). Preliminary work from our Lab also suggests this NT paradigm can improve spatial cognition, especially in experimental conditions associated with altered LTP/LTD ratios like aging and epilepsy (Amaro-Leal et al., 2016; Cunha-Reis, 2020). This suggests that this mismatch novelty training program may induce persistent changes at either synaptic domains or hippocampal circuits, favouring those altered responses to LTP or LTD/depotentiation induced by TBS or LFS. Possible mechanisms may involve altered physiology of hippocampal VIP expressing interneurons, that are targeted by many extrahippocampal projections regulating cognition during exploration and novelty/mismatch novelty responses. In fact, a subpopulation of VIP expressing interneurons recruited during theta oscillations (Luo et al., 2020) may play a role in information gating during spatial navigation and memory encoding. These are targets of *medium raphe* serotonergic and septal cholinergic and GABAergic projection fibres, that regulate the pacing, initiation, and suppressing of hippocampal theta rhythm (Borhegyi et al., 2004; Vandecasteele et al., 2014; Vinogradova et al., 1999). Furthermore, they are also targeted by prefrontal cortex long-range GABAergic projections (Malik et al., 2022), implicated in triggering hippocampal theta oscillations in response to mismatch computations (Garrido et al., 2015). As such, VIP expressing interneurons are in a key position to regulate many aspects of hippocampal dependent cognition related to mismatch novelty detection and processing.

VIP-mediated hippocampal disinhibition, being crucial for goal-directed spatial learning tasks (Turi et al., 2019) while suppressed to allow the hippocampal representation of objects during spatial exploration (Malik et al., 2022) may be differentially regulated by distinct types of novelty, known to trigger different levels of arousal and to selectively enhance activation of the hippocampus by serotonergic, noradrenergic, cholinergic and dopaminergic inputs (Walling et al., 2011; Schiffer et al., 2012; Schomaker and Meeter, 2015; Hagena and Manahan-Vaughan, 2017; Moreno-Castilla et al., 2017). These pathways are also involved in learning from prediction errors in humans (Schiffer et al., 2012), and differentially enhance/inhibit hippocampal synaptic plasticity in response to multiple behavioural stimuli (Hagena and Manahan-Vaughan, 2017). VIP immunoreactive interneurons control both feedforward and feedback inhibition being in a position to effectively modulate bidirectional hippocampal synaptic plasticity (Cunha-Reis and Caulino-Rocha, 2020). This, together with our recent findings that VIP VPAC_1_ receptor blockade could simultaneously enhance both hippocampal LTP and depotentiation (Cunha-Reis et al., 2014; Caulino-Rocha et al., 2022), led us to hypothesize that mechanisms controlled by VIP expressing interneurons could mediate, at least in part, the enhancing effects of mismatch novelty training on hippocampal LTP and depotentiation (Aidil-Carvalho et al., 2017). The observations in this work that mismatch novelty training does not change VIP or VPAC_1_ receptor levels in hippocampal synaptosomes, and that VPAC_1_ receptor mediated modulation of both LTP and depotentiation is impaired by mismatch novelty training appear to support this view. In particular, repeated inhibition or activation of distinct VIP-immunoreactive interneurons by mismatch novelty could favor an adaptation of synaptic communication and a decrease in the threshold for LTP and depotentiation induction. The influence on LTP is not likely mediated by a particular VIP-expressing interneuron subtype described to date only in mice, VIP long-range projection interneurons, that was reported to be active only when the hippocampal theta rhythm is supressed (Francavilla et al., 2018), as happening during attention shifts caused by novelty detection (Kitchigina et al., 1999). If present in the rat this may be responsible for endogenous VIP modulation of depotentiation.

It could be argued that VPAC_1_ receptors, also activated by PACAP and present in glutamatergic fibers in the hippocampus, could be the endogenous mediator of this effect. Yet, all the endogenous effects of PACAP reported so far in the hippocampus are mediated by either PAC_1_ or VPAC_2_ receptors (Cunha-Reis and Caulino-Rocha, 2020; Solés-Tarrés et al., 2020; Schmidt et al., 2021) and no dependency on GABAergic transmission has ever been attributed to those effects, making it very unlikely that PACAP the endogenous mediator of the effects here described. Altogether, the observations in this paper are consistent with our previous findings that basal synaptic transmission is decreased in NT animals (Aidil-Carvalho et al., 2017), that both LTP and depotentiation are enhanced in NT animals, and previous findings that mismatch novelty favors LTD and depotentiation over LTP induction *in vivo* (Kemp and Manahan-Vaughan, 2004; Sachser et al., 2017). This may be of particular relevance since depotentiation *in vivo* may be crucial for memory reformulation following mismatch detection (Goh and Manahan-Vaughan, 2013a).

Although it could be argued that the impaired effects of the VPAC_1_ antagonist on LTP/depotentiation in NT animals could be due to reaching a training-induced ceiling effect on LTP/depotentiation expression, given the mild TBS stimulation used to induce a moderate LTP, and the mild depression of synaptic transmission induced by the LFS protocol, it is not likely that the reduced effects of the VPAC_1_ antagonist on LTP were in fact limited by such a ceiling effect. Furthermore, as discussed above, the possibility that mismatch novelty training might have induced an adaptation in the activity of VIP immunoreactive interneurons or their interneuron targets is also not to be excluded and should be further investigated. Furthermore, a few observations in this study relate to the endogenous VIP modulation of depotentiation, that was abolished for HT and FT animal training and impaired in NT animals, even though depotentiation itself is enhanced in NT animals. Given the small effects (3-8%) involved in these differences, one should avoid the overinterpretation of these results. Nevertheless, one may argue that mismatch novelty stimuli are important in maintaining behavioural flexibility during learning, as opposed to repeated stimuli that not only keep depotentiation at its minimum, but also appear to abolish the possibility of its modulation by VIPergic interneurons.

The changes observed in VIP and VPAC_1_ receptor levels in HT and FT rats do not fully align with the observed effects on VPAC_1_ modulation of LTP and depotentiation. Being obtained from the full hippocampus and not only from the CA1 area, where changes in synaptic plasticity were studied, the sinaptossomal preparation used to quantify VIP and VPAC_1_ receptor levels may have masked the effects of mismatch novelty training on VIP or VPAC_1_ receptors in the CA1 area and the elucidation of this mechanism may require a different experimental approach. Regardless, the CA1 area of the hippocampus is known to be specifically activated by both spatial and temporal mismatch novelty in humans (Duncan et al., 2012; Chen et al., 2015; Thakral et al., 2015), and has for long been implicated in mismatch novelty detection and processing (Vinogradova, 2001).

Finally, even though changes in VPAC_1_ receptor mediated modulation of CA1 hippocampal plasticity by endogenous VIP may rely on subtle changes in disinhibition occurring only during LTP/depotentiation induction, we also showed that the training paradigms used in our work induced changes in the GABAergic synaptic content, as inferred from gephyrin synaptic immunoreactivity, and in the synaptic content of AMPA receptor GluA1 and GluA2 subunits and GluA1/GluA2 ratio, both very relevant for synaptic plasticity outcomes. Enhanced synaptic gephyrin immunoreactivity may either reflect an enhancement in hippocampal GABAergic synapses, that depending on the target could influence positively or negatively disinhibitory circuits, or may instead reflect enhanced gephyrin synaptic content overall in existing GABAergic synapses, which would lead to enhanced GABA_A_ receptor recruitment to postsynaptic sites (Pizzarelli et al., 2020). Interestingly, this enhancement was more pronounced in HT and FT animals than in NT animals, suggesting the enhancement is more related to the occurrence of daily training, than to mismatch novelty detection itself. Although altered levels of both GluA1, and most prominently GluA2 AMPA receptor subunits, were observed, the GluA1/GluA2 ratio was only significantly altered in NT animals. An enhanced GluA1/GluA2 ration may contribute to the enhanced LTP and depotentiation by NT by allowing a larger Ca^2+^ permeability (Chater and Goda, 2022), leaving synaptic plasticity at CA1 hippocampal synapses less susceptible to regulation by disinhibition control by endogenous VIP.

In conclusion, endogenous VIP, present on VIP immunoreactive interneurons, and acting on VPAC_1_ receptors, appears to be an important component of hippocampal-dependent mismatch novelty processing. The actions of VIPergic system can be altered by mismatch novelty training in a way that is consistent with the use of concurrent mechanisms to modulate CA1 hippocampal LTP and depotentiation. Although repetitive training, especially in the absence of objects, induced an enhancement of VPAC_1_ receptor expression this may not relate to functional changes in hippocampal CA1 area.

## Acknowledgements

D Cunha-Reis acknowledges Dr. Nadia C. Rodrigues for contribution in the setting up of original western blot experiments and the Institute of Physiology for animal housing facilities.

## Fellowships

SFRH/BPD/81358/2011 to DC-R and IPAD MSc Fellowship to FA-C.

## Researcher contract

Norma Transitória - DL57/2016/CP1479/CT0044 to DC-R. Funding sources made no contribution to the writing, research plan and decision to publish this paper.

## Author contributions

***F Aidil-Carvalho:*** formal analysis and methodology; ***A Caulino-Rocha:*** formal analysis and methodology; ***JA Ribeiro:*** supervision, funding acquisition, manuscript review and editing; ***D Cunha-Reis:*** Conceptualization, formal analysis and methodology, resources, supervision, funding acquisition, project administration, and writing – original draft, review, and editing.

## Data availability

The data that support the findings of this study are available from the corresponding author upon reasonable request.^1^

## Figure legends

**Supplementary Figure 1.**
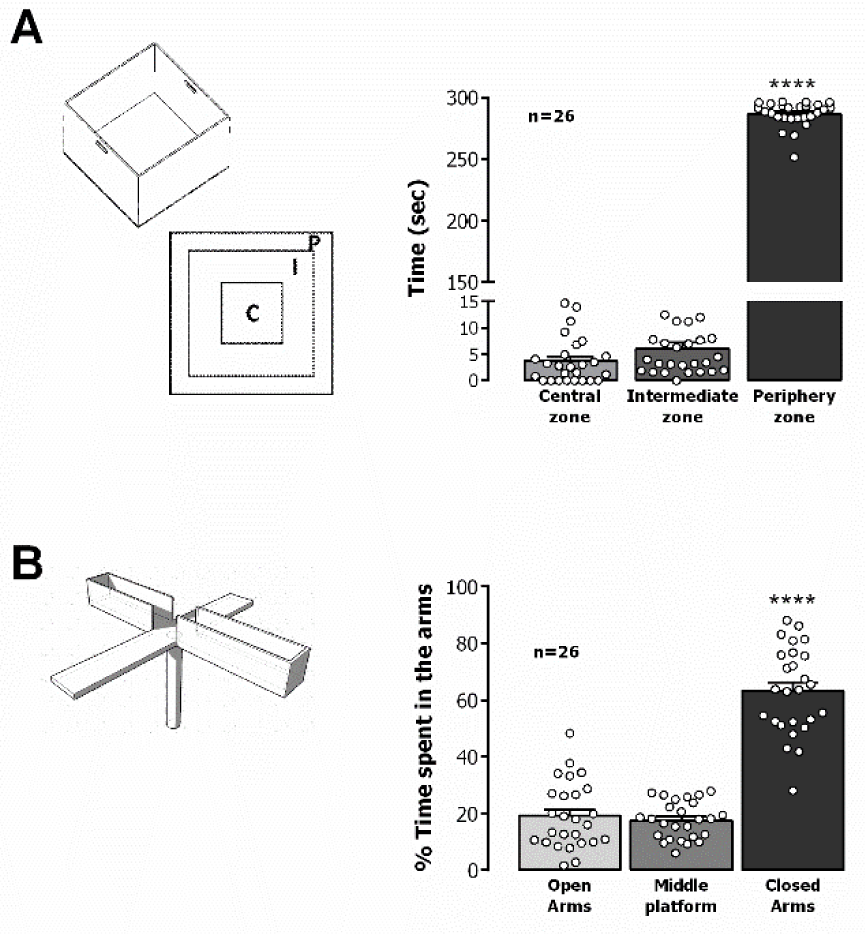
Behavioural evaluation prior to the mismatch novelty training paradigm. **A.** Evaluation of motor and exploratory capacity in the open-field (OF) test. ***p<0.005 (One-way ANOVA) as compared to central zone. **B.** Evaluation individual anxiety levels in the elevated plus maze (EPM) test. ****p<0.001 (One-way ANOVA) as compared to %time in open arms.

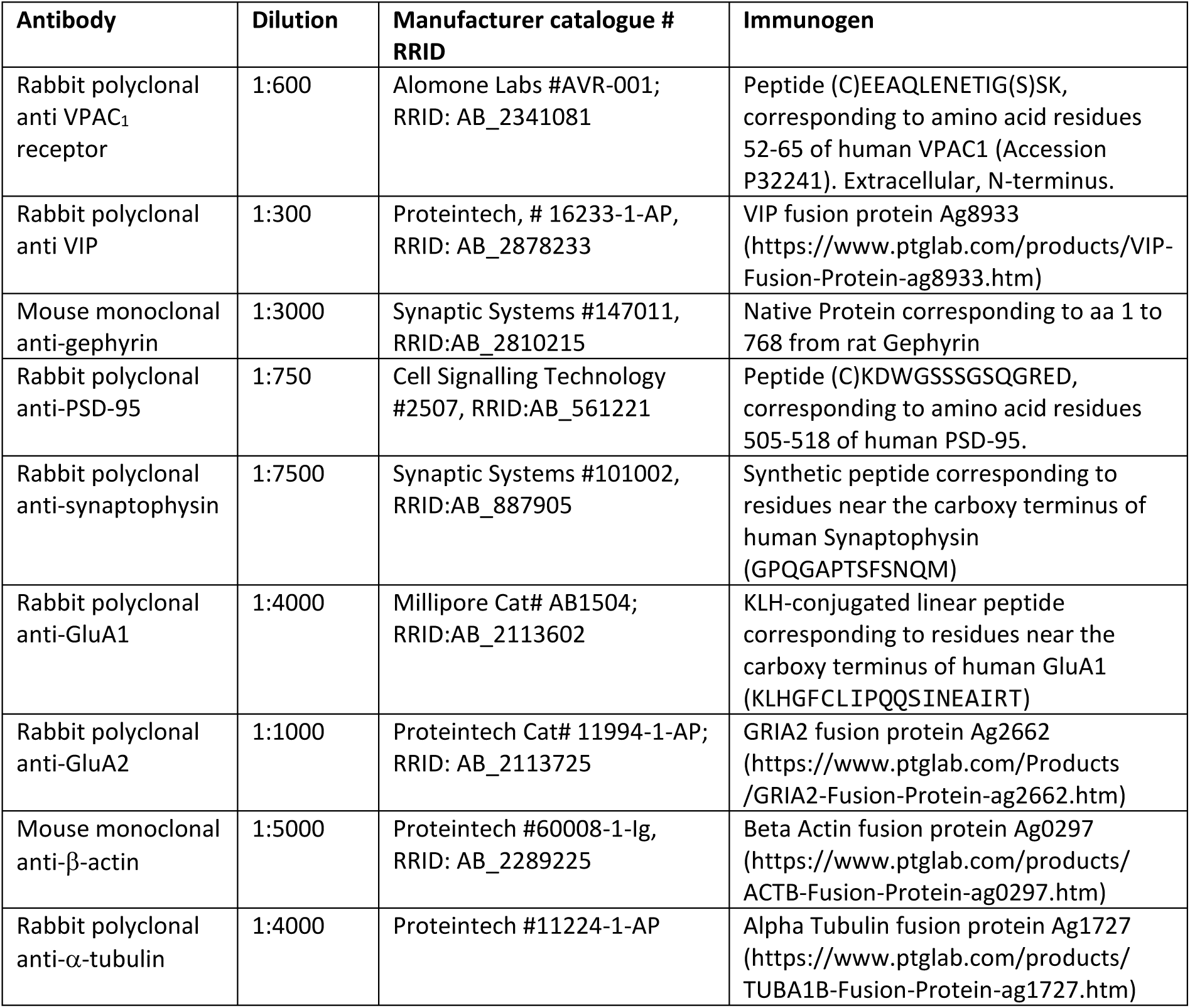
Antibody list.

## Footnotes

1 **Abbreviations:** aCSF, artificial cerebrospinal fluid; EPM, Elevated Plus Maze; FT, fixed training; HT, holeboard training; NT, mismatch novelty training; OF, Open-field; TBS, theta-burst stimulation.

## References

Abraham, W. C., and Bear, M. F. (1996). Metaplasticity: The plasticity of synaptic plasticity. Trends Neurosci. 19, 126–130. doi:10.1016/S0166-2236(96)80018-X.

Acsády, L., Görcs, T. J., and Freund, T. F. (1996). Different populations of vasoactive intestinal polypeptide-immunoreactive interneurons are specialized to control pyramidal cells or interneurons in the hippocampus. Neuroscience 73, 317–334. doi:10.1016/0306-4522(95)00609-5.

Aidil-Carvalho, M. F., Carmo, A. J. S., Ribeiro, J. A., and Cunha-Reis, D. (2017). Mismatch novelty exploration training enhances hippocampal synaptic plasticity: A tool for cognitive stimulation? Neurobiol. Learn. Mem. 145, 240–250. doi:10.1016/j.nlm.2017.09.004.

Alberini, C. M., and Ledoux, J. E. (2013). Memory reconsolidation. Curr. Biol. 23, R746–50. doi:10.1016/j.cub.2013.06.046.

Amaro-Leal, A., Rocha, I., and Cunha-Reis, D. (2016). Training in novelty exploration tasks prevents cognitive decline in a rat model of TLE. Epilepsia 57, 47.

Anderson, W. W., and Collingridge, G. L. (2001). The LTP Program: a data acquisition program for on-line analysis of long-term potentiation and other synaptic events. J Neurosci Methods 108, 71–83. doi:S0165-0270(01)00374-0 [pii].

Brown, G. R., and Nemes, C. (2008). The exploratory behaviour of rats in the hole-board apparatus: Is head-dipping a valid measure of neophilia? Behav. Processes 78, 442–448. doi:10.1016/j.beproc.2008.02.019.

Caulino-Rocha, A., Rodrigues, N. C., Ribeiro, J. A., and Cunha-Reis, D. (2022). Endogenous VIP VPAC1 Receptor Activation Modulates Hippocampal Theta Burst Induced LTP: Transduction Pathways and GABAergic Mechanisms. Biol. 2022, Vol. 11, Page 627 11, 627. doi:10.3390/BIOLOGY11050627.

Chater, T. E., and Goda, Y. (2022). The Shaping of AMPA Receptor Surface Distribution by Neuronal Activity. Front. Synaptic Neurosci. 14, 13. doi:10.3389/fnsyn.2022.833782.

Chen, J., Cook, P. A., and Wagner, A. D. (2015). Prediction strength modulates responses in human area CA1 to sequence violations. J. Neurophysiol. 114, 1227–1238. doi:10.1152/jn.00149.2015.

Crusio, W. E., Schwegler, H., and van Abeelen, J. H. F. (1989). Behavioral responses to novelty and structural variation of the hippocampus in mice. II. Multivariate genetic analysis. Behav. Brain Res. 32, 81–88. doi:10.1016/S0166-4328(89)80075-0.

Cunha-Reis, D. (2020). Mismatch Novelty Exploration Training Shaping Of Hippocampal Synaptic Plasticity And Cognition And The Role Of Disinhibition And VIP Expressing Interneurons. Biomed Biopharm Res 17, 3. doi:10.19277/bbr.17.2.243.

Cunha-Reis, D., Aidil-Carvalho, F., Ribeiro, J. A., Aidil-Carvalho, M. de F., and Ribeiro, J. A. (2014). Endogenous inhibition of hippocampal LTD and depotentiation by vasoactive intestinal peptide VPAC1 receptors. Hippocampus 24, 1353–1363. doi:10.1002/hipo.22316.

Cunha-Reis, D., and Caulino-Rocha, A. (2020). VIP Modulation of Hippocampal Synaptic Plasticity: A Role for VIP Receptors as Therapeutic Targets in Cognitive Decline and Mesial Temporal Lobe Epilepsy. *Front*. Cell. Neurosci. 14, 153. doi:10.3389/fncel.2020.00153.

Cunha-Reis, D., Caulino-Rocha, A., and Correia-de-Sá, P. (2021). VIPergic neuroprotection in epileptogenesis: challenges and opportunities. Pharmacol. Res. 164, 105356. doi:10.1016/j.phrs.2020.105356.

Cunha-Reis, D., Ribeiro, J. A., de Almeida, R. F. M., and Sebastião, A. M. (2017). VPAC1 and VPAC2 receptor activation on GABA release from hippocampal nerve terminals involve several different signalling pathways. Br. J. Pharmacol. 174, 4725–4737. doi:10.1111/bph.14051.

Dong, Z., Bai, Y., Wu, X., Li, H., Gong, B., Howland, J. G., et al. (2013). Hippocampal long-term depression mediates spatial reversal learning in the Morris water maze. Neuropharmacology 64, 65–73. doi:10.1016/j.neuropharm.2012.06.027.

Dong, Z., Gong, B., Li, H., Bai, Y., Wu, X., Huang, Y., et al. (2012). Mechanisms of hippocampal long-term depression are required for memory enhancement by novelty exploration. J. Neurosci. 32, 11980–11990. doi:10.1523/JNEUROSCI.0984-12.2012.

Duncan, K., Ketz, N., Inati, S. J., and Davachi, L. (2012). Evidence for area CA1 as a match/mismatch detector: A high-resolution fMRI study of the human hippocampus. Hippocampus 22, 389–398. doi:10.1002/hipo.20933.

Francavilla, R., Villette, V., Luo, X., Chamberland, S., Muñoz-Pino, E., Camiré, O., et al. (2018). Connectivity and network state-dependent recruitment of long-range VIP-GABAergic neurons in the mouse hippocampus. Nat. Commun. 9. doi:10.1038/s41467-018-07162-5.

Garrido, M. I., Barnes, G. R., Kumaran, D., Maguire, E. A., and Dolan, R. J. (2015). Ventromedial prefrontal cortex drives hippocampal theta oscillations induced by mismatch computations. Neuroimage 120, 362–370. doi:10.1016/j.neuroimage.2015.07.016.

Ge, Y., Dong, Z., Bagot, R. C., Howland, J. G., Phillips, A. G., Wong, T. P., et al. (2010). Hippocampal long-term depression is required for the consolidation of spatial memory. Proc. Natl. Acad. Sci. U. S. A. 107, 16697–16702. doi:10.1073/pnas.1008200107.

Goh, J. J., and Manahan-Vaughan, D. (2013a). Hippocampal long-term depression in freely behaving mice requires the activation of beta-adrenergic receptors. Hippocampus 23, 1299–1308. doi:10.1002/hipo.22168.

Goh, J. J., and Manahan-Vaughan, D. (2013b). Spatial object recognition enables endogenous LTD that curtails LTP in the mouse hippocampus. Cereb. Cortex 23, 1118–25. doi:10.1093/cercor/bhs089.

Hagena, H., and Manahan-Vaughan, D. (2017). The serotonergic 5-HT4 receptor: A unique modulator of hippocampal synaptic information processing and cognition. Neurobiol. Learn. Mem. 138, 145–153. doi:10.1016/j.nlm.2016.06.014.

Izquierdo, L. A., Viola, H., Barros, D. M., Alonso, M., Vianna, M. R. M., Furman, M., et al. (2001). Novelty enhances retrieval: Molecular mechanisms involved in rat hippocampus. Eur. J. Neurosci. 13, 1464–1467. doi:10.1046/j.0953-816X.2001.01530.x.

Kanari, K., Kikusui, T., Takeuchi, Y., and Mori, Y. (2005). Multidimensional structure of anxiety-related behavior in early-weaned rats. Behav. Brain Res. 156, 45–52. doi:10.1016/j.bbr.2004.05.008.

Kemp, A., and Manahan-Vaughan, D. (2004). Hippocampal long-term depression and long-term potentiation encode different aspects of novelty acquisition. Proc Natl Acad Sci U S A 101, 8192–8197. doi:0402650101 [pii] 10.1073/pnas.0402650101.

Kitchigina, V. F., Kudina, T. A., Kutyreva, E. V, and Vinogradova, O. S. (1999). Neuronal activity of the septal pacemaker of theta rhythm under the influence of stimulation and blockade of the median raphe nucleus in the awake rabbit. Neuroscience 94, 453–463. doi:10.1016/s0306-4522(99)00258-4.

Luo, X., Guet-Mccreight, A., Villette, V., Francavilla, R., Marino, B., Chamberland, S., et al. (2020). Synaptic Mechanisms Underlying the Network State-Dependent Recruitment of VIP-Expressing Interneurons in the CA1 Hippocampus. Cereb. Cortex 30, 3667–3685. doi:10.1093/cercor/bhz334.

Malik, R., Li, Y., Schamiloglu, S., and Sohal, V. S. (2022). Top-down control of hippocampal signal-to-noise by prefrontal long-range inhibition. Cell 185, 1602–1617.e17. doi:10.1016/j.cell.2022.04.001.

Manahan-Vaughan, D., and Braunewell, K. H. (1999). Novelty acquisition is associated with induction of hippocampal long-term depression. Proc Natl Acad Sci U S A 96, 8739–8744. Available at: http://www.ncbi.nlm.nih.gov/pubmed/10411945.

Moncada, D., and Viola, H. (2007). Induction of long-term memory by exposure to novelty requires protein synthesis: Evidence for a behavioral tagging. J. Neurosci. 27, 7476–7481. doi:10.1523/JNEUROSCI.1083-07.2007.

Moreno-Castilla, P., Pérez-Ortega, R., Violante-Soria, V., Balderas, I., and Bermúdez-Rattoni, F. (2017). Hippocampal release of dopamine and norepinephrine encodes novel contextual information. Hippocampus 27, 547–557. doi:10.1002/hipo.22711.

Park, A. J., Harris, A. Z., Martyniuk, K. M., Chang, C. Y., Abbas, A. I., Lowes, D. C., et al. (2021). Reset of hippocampal–prefrontal circuitry facilitates learning. Nature 591, 615–619. doi:10.1038/s41586-021-03272-1.

Pedreira, M. E., Pérez-Cuesta, L. M., and Maldonado, H. (2004). Mismatch between what is expected and what actually occurs triggers memory reconsolidation or extinction. Learn. Mem. 11, 579–585. doi:10.1101/lm.76904.

Pizzarelli, R., Griguoli, M., Zacchi, P., Petrini, E. M., Barberis, A., Cattaneo, A., et al. (2020). Tuning GABAergic Inhibition: Gephyrin Molecular Organization and Functions. Neuroscience 439, 125–136.

Qi, Y., Hu, N. W., and Rowan, M. J. (2013). Switching off LTP: MGlu and NMDA receptor-dependent novelty exploration-induced depotentiation in the rat hippocampus. Cereb. Cortex 23, 932–939. doi:10.1093/cercor/bhs086.

Quent, J. A., Henson, R. N., and Greve, A. (2021). A predictive account of how novelty influences declarative memory. Neurobiol. Learn. Mem. 179, 107382. doi:10.1016/j.nlm.2021.107382.

Ribeiro, J. A., Cunha-Reis, D., Lopes, L. V., Coelho, J. E., Costenla, A. R., Correia-de-Sá, P., et al. (2001). Adenosine receptor interactions in the hippocampus. Drug Dev. Res. 52, 337–345. doi:10.1002/ddr.1132.

Rodrigues, N. C., Silva-Cruz, A., Caulino-Rocha, A., Bento-Oliveira, A., Alexandre Ribeiro, J., and Cunha-Reis, D. (2021). Hippocampal CA1 theta burst-induced LTP from weaning to adulthood: Cellular and molecular mechanisms in young male rats revisited. Eur. J. Neurosci. 54, 5272–5292. doi:10.1111/ejn.15390.

Sachser, R. M., Haubrich, J., Lunardi, P. S., and de Oliveira Alvares, L. (2017). Forgetting of what was once learned: Exploring the role of postsynaptic ionotropic glutamate receptors on memory formation, maintenance, and decay. Neuropharmacology 112, 94–103. Available at: http://www.sciencedirect.com/science/article/pii/S002839081630301X?via%3Dihub [Accessed July 29, 2017].

Schiffer, A. M., Ahlheim, C., Wurm, M. F., and Schubotz, R. I. (2012). Surprised at all the entropy: Hippocampal, caudate and midbrain contributions to learning from prediction errors. PLoS One 7, e36445. doi:10.1371/journal.pone.0036445.

Schmidt, S. D., Zinn, C. G., Behling, J. A. K., Furian, A. F., Furini, C. R. G., de Carvalho Myskiw, J., et al. (2021). Inhibition of PACAP/PAC1/VPAC2 signaling impairs the consolidation of social recognition memory and nitric oxide prevents this deficit. Neurobiol. Learn. Mem. 180, 107423. doi:10.1016/J.NLM.2021.107423.

Schneider, P., Ho, Y.-J., Spanagel, R., and Pawlak, C. R. (2011). A Novel Elevated Plus-Maze Procedure to Avoid the One-Trial Tolerance Problem. Front. Behav. Neurosci. 5, 43. doi:10.3389/fnbeh.2011.00043.

Schomaker, J., and Meeter, M. (2015). Short- and long-lasting consequences of novelty, deviance and surprise on brain and cognition. Neurosci Biobehav Rev 55, 268–279. doi:10.1016/j.neubiorev.2015.05.002.

Solés-Tarrés, I., Cabezas-Llobet, N., Vaudry, D., and Xifró, X. (2020). Protective Effects of Pituitary Adenylate Cyclase-Activating Polypeptide and Vasoactive Intestinal Peptide Against Cognitive Decline in Neurodegenerative Diseases. Front. Cell. Neurosci. 14, 221. doi:10.3389/fncel.2020.00221.

Thakral, P. P., Yu, S. S., and Rugg, M. D. (2015). The hippocampus is sensitive to the mismatch in novelty between items and their contexts. Brain Res 1602, 144–152. doi:10.1016/j.brainres.2015.01.033.

Trempler, I., Schiffer, A. M., El-Sourani, N., Ahlheim, C., Fink, G. R., and Schubotz, R. I. (2017). Frontostriatal contribution to the interplay of flexibility and stability in serial prediction. J. Cogn. Neurosci. 29, 298–309. doi:10.1162/jocn_a_01040.

Turi, G. F., Li, W. K., Chavlis, S., Pandi, I., O’Hare, J., Priestley, J. B., et al. (2019). Vasoactive Intestinal Polypeptide-Expressing Interneurons in the Hippocampus Support Goal-Oriented Spatial Learning. Neuron 101, 1150–1165.e8. doi:10.1016/j.neuron.2019.01.009.

Vinogradova, O. S. S. (2001). Hippocampus as comparator: Role of the two input and two output systems of the hippocampus in selection and registration of information. Hippocampus 11, 578–598. doi:10.1002/hipo.1073.

Wagner, J. J., and Alger, B. E. (1995). GABAergic and developmental influences on homosynaptic LTD and depotentiation in rat hippocampus. J Neurosci 15, 1577–1586. Available at: http://www.ncbi.nlm.nih.gov/pubmed/7869119.

Walling, S. G., Brown, R. A. M., Milway, J. S., Earle, A. G., and Harley, C. W. (2011). Selective tuning of hippocampal oscillations by phasic locus coeruleus activation in awake male rats. Hippocampus 21, 1250– 1262. doi:10.1002/hipo.20816.

